# Aversion of the invasive Asian longhorned tick to the white-footed mouse, the dominant reservoir of tick-borne pathogens in the United States

**DOI:** 10.1101/826289

**Authors:** Isobel Ronai, Danielle M. Tufts, Maria A. Diuk-Wasser

**Affiliations:** Department of Ecology, Evolution, & Environmental Biology, Columbia University

**Keywords:** *Canis lupus familiaris*, *Felis catus*, *Homo sapiens*, *Odocoileus virginianus*, acquired tick resistance, blacklegged tick, host immunity, host-seeking, hot foot, Ixodidae

## Abstract

The Asian longhorned tick (*Haemaphysalis longicornis*) was reported for the first time in the United States of America in 2017 and has now spread across 12 states. The potential of this invasive tick vector to transmit pathogens will be determined through its association to native hosts, such as the white-footed mouse (*Peromyscus leucopus*) which is the primary reservoir for the causative agent of Lyme disease (*Borrelia burgdorferi*) and other zoonotic pathogens. We placed larval *H. longicornis* on *P. leucopus*, 65% of the larvae (*n* = 40) moved off the host within a short period of time and none engorged. In contrast, larval black-legged ticks (*Ixodes scapularis*) did not move from where they were placed in the ear of the mouse. We then conducted a laboratory behavioural assay to assess the interaction of *H. longicornis* with the hair of potential mammalian host species in the United States of America. *H. longicornis* larvae were less likely to enter the hair zone of *P. leucopus* and humans compared to the hair of domestic cats, domestic dogs, and white-tailed deer. Our study identifies a tick-host hair interaction behaviour, which can be quantified in a laboratory assay to predict tick-host associations and provides insights into how ticks select a host.

## Introduction

The Asian longhorned tick (*Haemaphysalis longicornis*) transmits numerous human pathogens and is a highly invasive tick species [1-3]. In the United States of America this species was reported for the first time in 2017 [4], although archival evidence suggests *H. longicornis* has been present in the USA since 2010 [5]. Currently, *H. longicornis* has been detected in 12 states: Arkansas, Connecticut, Delaware, Kentucky, Maryland, New Jersey, New York, North Carolina, Pennsylvania, Tennessee, Virginia and West Virginia [2, 6]. Modelling studies indicate this species has the potential to spread throughout the majority of the USA [7].

As *H. longicornis* establishes and spreads to new ecosystems it encounters new host communities. The host blood meal is critical for not only vector survival and reproduction but also for the pathogens it can acquire and transmit. In the USA, the non-domestic mammalian host community for ticks includes: small mammals (such as white-footed mice and other rodents); medium mammals (such as racoons and opossum); and large mammals (such as deer) [8]. The host species of most importance for public health is the white-footed mouse (*Peromyscus leucopus*), the primary vertebrate reservoir host for zoonotic pathogens, such as the causative agent of Lyme disease (*Borrelia burgdorferi*) [8]. However, larval *H. longicornis* have shown limited association with small rodents compared to medium and large sized mammals, in both its native and invasive ranges [1, 3, 9, 10].

Here we investigate the interaction of the invasive *H. longicornis* larvae with *P. leucopus* and other potential mammalian host species commonly encountered in the USA, including humans. The behaviour of *H. longicornis* is also compared to that of the native black-legged tick (*Ixodes scapularis*), the main vector of *B. burgdorferi* and at least six other human pathogens in the USA [11].

## Materials and Methods

### Ticks

During fieldwork on Staten Island (New York, USA) in August 2018, we collected engorged *H. longicornis* adult females from a white-tailed deer (*Odocoileus virginianus*) [10]. A subset of these adult females were tested for pathogens and found to be negative. Three females were maintained in individual vials in an incubator (21°C add 95-100% humidity, light:dark cycle was 16:8 h to simulate summer conditions) and allowed to lay eggs. Larvae emerged from the egg masses 4 months later. The *I. scapularis* larvae were obtained from a laboratory-reared colony through the NIH Biodefense and Emerging Infections Research Resources Repository (NIAID, NIH: *I. scapularis* larvae, NR-44115). These larvae were maintained in the same incubator and used in the study within 6 months.

### Behavioural assessment of responses to live white-footed mouse host

We placed 10 *H. longicornis* (*n* = 4 replicates) or 10 *I. scapularis* larvae (*n* = 3 replicates) in the ear canal of an anaesthetised mouse (*n* = 4, 2 replicates per mouse, one per ear). The behaviour of the ticks in the ear canal of the anaesthetised mouse was observed every 30 seconds for 15 minutes and we noted the duration of time the ticks took to: (i) move off the ear of the mouse; and (ii) drop off the mouse. To investigate whether the remaining *H. longicornis* would feed to repletion they were left on the mice. Individual mice were housed in single cages positioned over water. The mouse cages were inspected daily for any engorged larvae and the number of recovered larvae were recorded. All animal procedures were in accordance with guidelines approved by the Columbia University Institutional Animal Care and Use Committee (IACUC), protocol number: AC-AAAY2450.

### Behavioural arena assay of interaction with potential native hosts

Hair was removed from: three frozen white-footed mice (*P. leucopus*); two domestic cats (*Felis catus*); one domestic dog (*Canis lupus familiaris*); multiple white-tailed deer; and one human (*Homo sapiens*). None of the animals were treated with flea or tick repellent. The human hair was obtained from the head of one of the researchers (DMT) and was not dyed or treated with any chemicals. Petri dishes (150 mm x 15 mm) were used as behavioural arenas and the dish was divided into two areas (the hair zone and non-hair zone divided by a centre line) (figure 1). The hair was arranged in a layer inside the hair zone and a new Petri dish was used for each hair treatment to prevent scent cross-contamination. At time zero, we placed *H. longicornis* or *I. scapularis* larvae (*n* = 10, 3 replicates per tick species per hair treatment) on the centre line. Any tick that moved to the rim of the Petri dish was relocated with a brush to the base of the dish directly below the rim.

**Figure 1.**
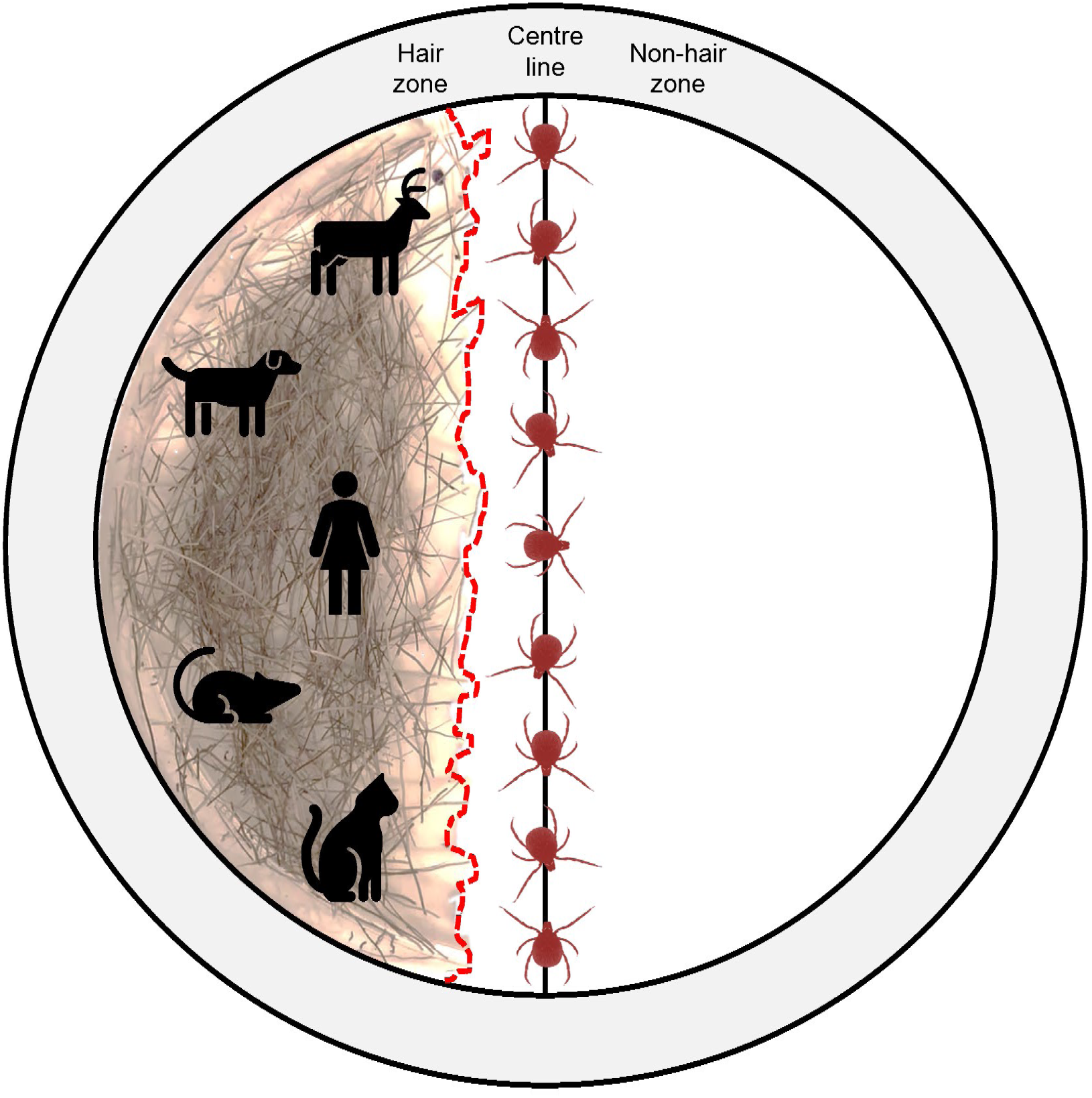
Diagram of behavioural assay arena. Petri dish (150 mm × 15 mm) divided into three zones: centre line, hair zone, and non-hair zone. The host hair (white-footed mouse, cat, dog, white-tailed deer, and human) was placed in the hair zone and formed an irregular hair interface (dashed line). At the start of the behavioural assay *Haemaphysalis longicornis* or *Ixodes scapularis* larvae (*n* = 10) were placed on the centre line, assays were replicated three times for each hair treatment.

To assess the behaviour of the ticks when encountering host hair, each trial of the behavioural assay was video recorded for 5 minutes. The videos were analysed by two double-blinded observers. The main behavioural response of the ticks was an interaction with the hair interface (come within 1 mm of the hair or touch the hair) which is the edge of the hair zone (dotted line in figure 1). We counted the number of times the ticks interacted with the hair interface and report the frequency of interactions per tick per minute. Note that a tick sometimes interacted with the hair interface multiple times. We then recorded the outcome of the interaction, the tick either: (i) entered the hair zone (supporting information video 1); or (ii) turned away from the hair interface (supporting information video 2).

### Statistical analysis

We conducted all statistical analyses using R software. The effect of tick species and host hair treatment on the number of times ticks interacted with the hair interface was examined using a non-parametric Kruskal-Wallis test. The resulting decision (entered the hair zone or turned away at the hair interface) given an interaction was assessed using a generalised linear mixed model (GLMM) fit by maximum likelihood (Laplace approximation) with observer and replicate as random effects (R package lme4). A GLMM model was performed to compare the interaction behaviour between the two species of tick and then separate analyses were performed for each tick species to compare the probability of entering the hair zone of different hosts to that of the white-footed mouse (reference category). Lastly, a student’s t-test was used to compare the time individual ticks spent in the hair zone comparing *H. longicornis* and *I. scapularis*.

## Results

Within a 15-minute timeframe after placement on white-footed mice, 67.5% of the *H. longicornis* (*n* = 40) moved from the site of placement inside the mouse ear canal, whereas 0% of the *I. scapularis* (*n* = 40) moved (supporting information, figure S1a). In addition, 55% of the *H. longicornis* (*n* = 40) dropped off the mice, whereas 0% of *I. scapularis* (*n* = 40) dropped off (supporting information, figure S1b). Before we relocated the mice to a cage, an additional four *H. longicornis* dropped off, therefore, 65% of *H. longicornis* (*n* = 40) dropped off the mice. No engorged larvae of the remaining *H. longicornis* on the mice (*n* = 14) were recovered.

*H. longicornis* and *I. scapularis* had a similar frequency of interactions with all of the hair treatment interfaces (Kruskal-Wallis: 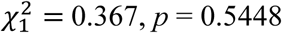; supporting information, figure S2). There was also no significant effect of hair treatments across the two species of tick (Kruskal-Wallis: 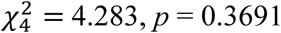; supporting information, figure 2). Therefore, *H. longicornis* interacted as frequently with the hair treatments as *I. scapularis*.

**Figure 2.**
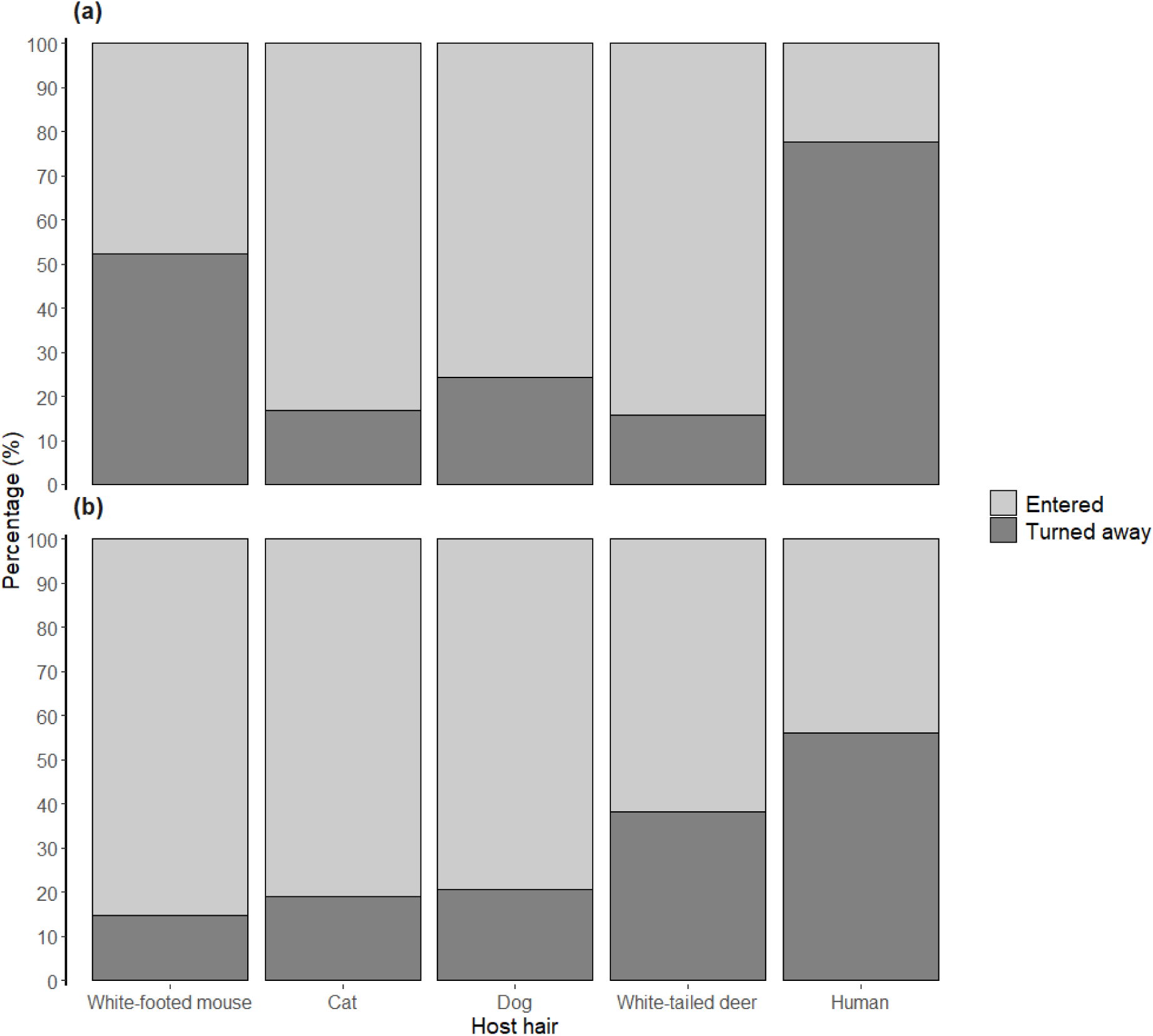
The percentage of ticks that, on interaction with the interface of the host hair (white-footed mouse, cat, dog, white-tailed deer, and human), either entered the hair zone or turned away from the interface for (a) *Haemaphysalis longicornis* larvae and (b) *Ixodes scapularis* larvae.

We observed that when a tick interacted with the hair interface they raised their front legs (the location of the sensory Haller’s organ [12]) and waved them (supporting information videos 1 & 2). After each interaction, the tick decided to either enter the host hair zone or turn away from the hair interface. We used this decision by the ticks as a behavioural metric.

After an interaction with the hair interface *H. longicornis* larvae were significantly less likely to enter the host hair zone compared to *I. scapularis* larvae (GLMM, *p* = 0.0365, figure 2, table 1). We then analysed the behaviour within each tick species. *H. longicornis* larvae were significantly more likely to enter the hair zone of cats, dogs, or white-tailed deer than the hair zone of white-footed mice (*p* = 0.0095; *p* = 0.0261; and *p* = 0.0039; respectively, figure 2a, table 1). In addition, *H. longicornis* larvae were as likely to enter the hair zone of humans as the hair zone of white-footed mice (*p* = 0.1645, figure 2a, table 1). *I. scapularis* larvae were significantly more likely to enter the hair zone of white-footed mice than the hair zone of white-tailed deer, or humans (*p* = 0.0447; and *p* = 0.0021; respectively, figure 2b, table 1). In addition, *I. scapularis* larvae were as likely to enter the hair zone of white-footed mice as the hair zone of cats or dogs (*p* = 0.3415; and *p* = 0.4094; respectively, figure 2b, table 1). We also found that *H. longicornis* larvae spent significantly less time within the hair zone of each hair treatment compared to *I. scapularis* larvae (*p* = 0.0040).

**Table 1.** GLMM of *Haemaphysalis longicornis* larvae and *Ixodes scapularis* larvae. The number of individual larval interactions (*n*) with the host hair interface (white-footed mouse, cat, dog, white-tailed deer, and human), calculated odds ratio, and p-value. Each treatment was compared to the white-footed mouse (reference). (*) P < 0.05 and (**) P < 0.01.

| Treatment | *Haemaphysalis longicornis* |  |  | *Ixodes scapularis* |  |  |
| --- | --- | --- | --- | --- | --- | --- |
|  | *n* | Odds Ratio | *P*-value | *n* | Odds Ratio | *P*-value |
| White-footed mouse | 37 | 1 | - | 33 | 1 | - |
| Cat | 23 | 5.0901 | 0.0096** | 8 | 0.5537 | 0.3415 |
| Dog | 41 | 3.1535 | 0.0261* | 34 | 0.5915 | 0.4094 |
| White-tailed deer | 22 | 6.9331 | 0.0039** | 22 | 0.2830 | 0.0447* |
| Human | 29 | 0.4616 | 0.1645 | 25 | 0.1381 | 0.0021** |

## Discussion

We observed that host-seeking *H. longicornis* larvae exhibited aversion to the hair of white-footed mice. This newly invasive tick is therefore unlikely to select the white-footed mouse as a host in the natural environment of the USA. The findings of our laboratory-based study help explain why the recent USA passive and active field studies of *H. longicornis* did not find *H. longicornis* of any life stage on white-footed mice, despite collection of host-seeking *H. longicornis* ticks in the same regions [5, 10]. The aversion of *H. longicornis* to the white-footed mouse and humans means there is a lower likelihood for them to acquire and be a vector of the most important tick-borne pathogens (such as *B. burgdorferi, Babesia microti* and *Anaplasma phagocytophilum*) in the USA, for which white-footed mice are the main reservoir host [11]. Furthermore, *H. longicornis* were unable to maintain *B. burgdorferi* transstadially after feeding on infected *Mus musculus* while confined in feeding capsules [13].

Why has *H. longicornis* evolved an aversion to the white-footed mouse? A possible explanation for this aversion is that feeding on mice reduces the fitness of *H. longicornis*. For example, house mice develop resistance to *H. longicornis* after one exposure [14] which results in a reduction in fitness for the ticks (detachment from host, prolonged duration for feeding on a host, impaired engorgement, lower egg clutch sizes, and moulting death [15]). The acquired tick resistance of mice is due to an immunological response via antibody production [14, 15].

Our findings that larval *H. longicornis* are more likely to enter the hair zone of medium and large sized mammals is consistent with field studies of *H. longicornis* [5, 10]. Medium-sized mammals have intermediate competence for tick-borne pathogens such as *B. burgdorferi* [8] and it is currently unknown whether medium-sized mammals can serve as a source of pathogens for *H. longicornis* in the USA. In addition, we found larval *H. longicornis* have an aversion for human hair. Notably, there are only two cases so far reported of *H. longicornis* biting a human in the USA [5].

On physical contact with a passing potential host, a tick must make a ‘decision’ whether or not to hold onto the host and subsequently feed. How different tick species detect and then select their host is currently not well understood [12]. Host stimuli such as body heat and carbon dioxide are not species specific so likely unhelpful for host selection. Our study has identified that ticks have a unique hair interaction behaviour (decide to enter the host hair or turn away at the hair interface), which suggests that they utilise a species-specific property of the animal hair to select their host. Furthermore, a behavioural assay that utilises host hair could provide a measure of potential tick-host associations that do not yet occur in nature, such as newly invasive ticks or ticks expanding their geographic range.

In conclusion, our study finds that the newly invasive *H. longicornis* has an aversion to the white-footed mouse, the dominant reservoir of tick-borne pathogens in the USA. *H. longicornis* also has an aversion to humans which decreases human risk for tick bites. Pathogen transmission studies therefore need to consider not only attraction of a vector to a host but also host aversion.

## Acknowledgments

We wish to thank Kevin Zhao and Daniel Mathisson for assisting with video analysis, and Pilar Fernandez for advice on data analysis. In addition, thanks to Thomas Hart who provided the deer hair and the pets (Venus, Luna, and Lucy) who contributed their hair.

