## Supplementary material for "Aversion of the invasive Asian longhorned tick to the white-footed mouse, the dominant reservoir of tick-borne pathogens in the United States"

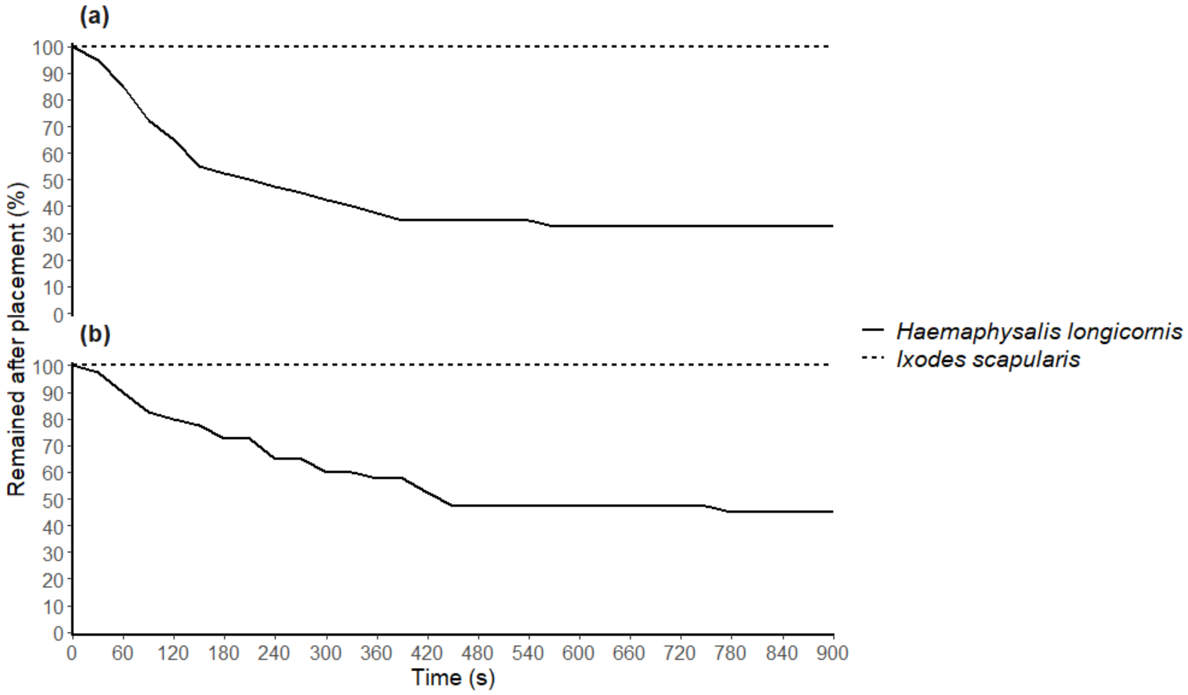


**Figure 1.** Percentage of *Haemaphysalis longicornis* (*n* = 40) or *Ixodes scapularis* (*n* = 30) larvae that remained after placement (time 0) on the (a) ear of the mouse and (b) mouse.

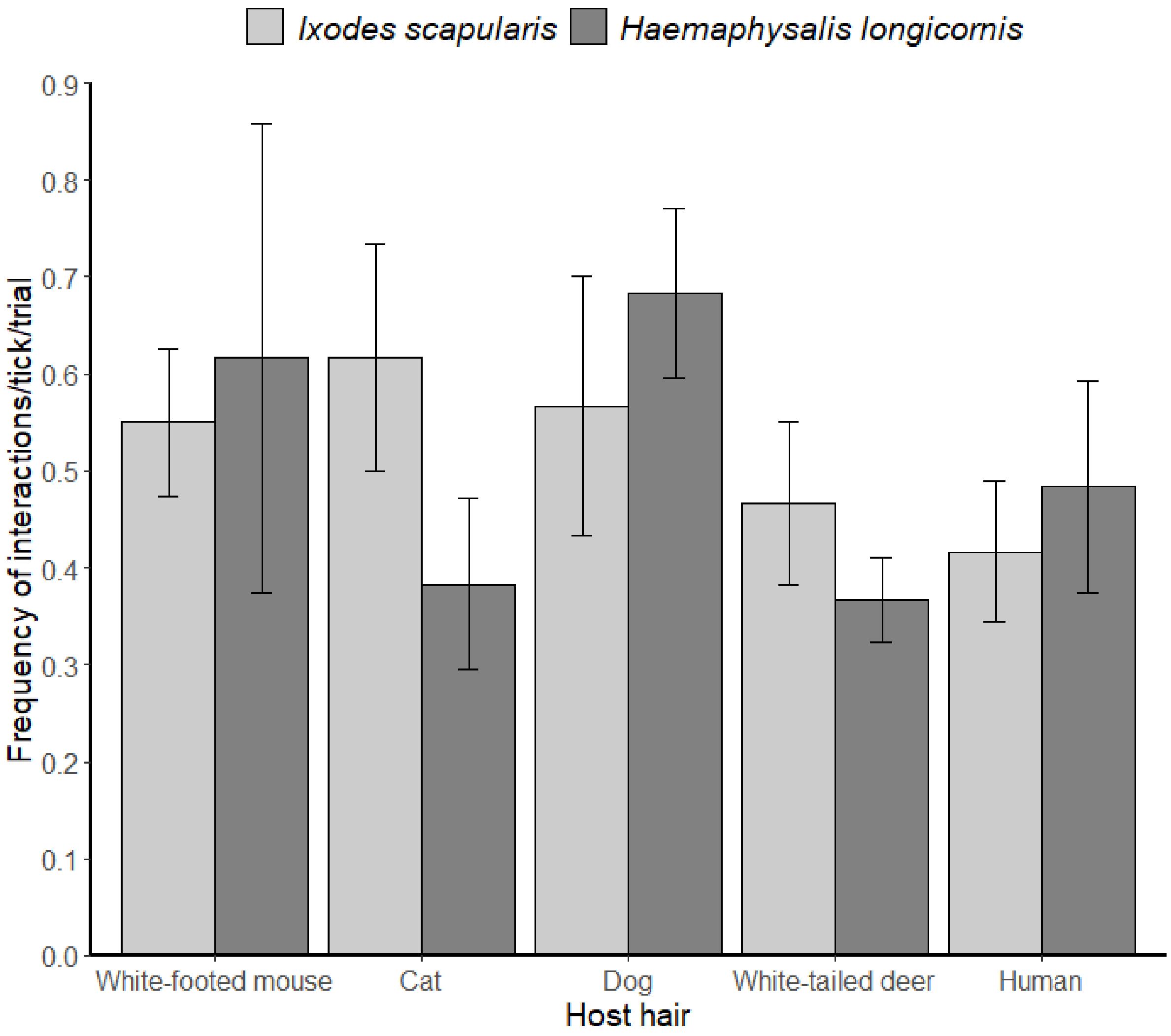


**Figure 2**. Number of interactions a larval tick (*Haemaphysalis longicornis* or *Ixodes scapularis*, *n* = 30) made with the interface of the host hair (white-footed mouse, cat, dog, white-tailed deer, and human) per trial.
